# Landscape of infection enhancing antibodies in COVID-19 and healthy donors

**DOI:** 10.1101/2022.07.09.499414

**Authors:** Hendra S Ismanto, Zichang Xu, Dianita S Saputri, Jan Wilamowski, Songling Li, Dendi K Nugraha, Yasuhiko Horiguchi, Masato Okada, Hisashi Arase, Daron M Standley

**Affiliations:** Department of Genome Informatics, Research Institute for Microbial Diseases, Osaka University, 3-1 Yamadaoka, Suita 565-0871, Japan; Department of Immunochemistry, Research Institute for Microbial Diseases, Osaka University, 3-1 Yamadaoka, Suita 565-0871, Japan; Deparment of Molecular Bacteriology, Research Institute for Microbial Diseases, Osaka University, 3-1 Yamadaoka, Suita 565-0871, Japan; Deparment of Oncogene Research, Research Institute for Microbial Diseases, Osaka University, 3-1 Yamadaoka, Suita 565-0871, Japan; Department of System Immunology, Immunology Frontier Research Center, Osaka University, 3-1 Yamadaoka, Suita 565-0871, Japan; Department of Immunochemistry, Immunology Frontier Research Center, Osaka University, 3-1 Yamadaoka, Suita 565-0871, Japan; Department of Oncogene Research, Immunology Frontier Research Center, Osaka University, 3-1 Yamadaoka, Suita 565-0871, Japan; Center for Infectious Disease Education and Research, Osaka University, Osaka 565-0871, Japan

**Keywords:** COVID-19, SARS-CoV-2, Infection enhancing antibodies, Antibody repertoire, InterClone

## Abstract

To assess the frequency of SARS-CoV-2 infection enhancing antibodies in the general population, we searched over 64 million heavy chain antibody sequences from healthy and COVID-19 patient repertoires for sequences similar to 11 previously reported enhancing antibodies. Although the distribution of sequence identities was similar in COVID-19 and healthy repertoires, the COVID-19 hits were significantly more clonally expanded than healthy hits. Furthermore, among the tested hits, 17 out of 94 from COVID-19, compared with 2 out of 96 from healthy, bound to the enhancing epitope. A total of 6 of the 19 epitope-binding antibodies enhanced ACE2 receptor binding to the spike protein. Together, this study revealed that enhancing antibodies are far more frequent in COVID-19 patients than in healthy donors, but a reservoir of potential enhancing antibodies exists in healthy donors that could potentially mature to actual enhancing antibodies upon infection.

## Introduction

Upon virus infection, host B cells that recognize viral antigens undergo affinity maturation and differentiation into antibody-producing cells and memory B cells (Harwood & Batista, 2010; Victora & Nussenzweig, 2012). Together with T cells, antigen-specific neutralizing antibodies resolve infection (Dorner & Radbruch, 2007; Morales-Nunez et al., 2021), while long-lived memory B cells protect against future infections (Akkaya et al., 2020; Cyster & Allen, 2019; Kurosaki et al., 2015; Phan & Tangye, 2017). In the process of producing neutralizing antibodies, infection-enhancing antibodies can also be generated (Bournazos et al., 2020). Antibody-dependent enhancement (ADE) has been observed for multiple virus infections (Guzman et al., 2013; Kapikian et al., 1969; Kim et al., 1969; Polack et al., 2003; Simmons et al., 2012) and represents a challenge for the design of safe and effective vaccines (Arvin et al., 2020; Haynes et al., 2020).

In 2021, two groups independently identified antibodies that enhanced SARS-CoV-2 spike protein binding to human ACE2 (Li et al., 2021; Liu et al., 2021). Interestingly, the 11 monoclonal antibodies collectively identified in these two studies were distinct in terms of their amino acid sequences and gene usage yet targeted an overlapping site on the N-terminal domain of the spike protein. Although the molecular mechanism of the observed ACE2-binding and infection enhancement has not been demonstrated conclusively, multiple lines evidence point to a model involving crosslinking of adjacent spike proteins. This evidence includes cell-based assays showing that enhancement did not depend on the Fc domain of the antibody but did require two Fab arms (i.e., full-length IgG or F(ab’)2), as well as molecular modelling that indicated that the two Fab arms could not reach two enhancing epitopes on a single spike.

Since the proposed SARS-CoV-2 infection enhancing mechanism appears to be distinct from previously reported ADE models, we sought to quantify the frequency of sequences similar to the known enhancing antibodies in healthy and COVID-19 donors. Based on known structural data, most antibodies recognize their cognate antigens through their complementarity-determining regions (CDRs). Moreover, cryo-EM structural models of 3 out of the 11 enhancing antibodies indicate that most of the physical contacts are mediated by the heavy chain (Li et al., 2021; Liu et al., 2021). We thus reasoned that potential infection-enhancing antibodies could be identified through similarity to heavy chain CDRs. To this end, we utilized a bioinformatics pipeline for identifying antibodies in large BCR repertoire datasets with similar CDR sequences to a set of queries (Figure 1). We also performed antibody expression and binding assays to assess the functional phenotype of these antibodies among our search hits. Although they were less frequent than in COVID-19 patients, we identified potential enhancing antibodies in healthy donors that could lead to the development of actual enhancing antibodies upon infection. This study illustrates that large BCR repertoire data can be used to discover functional human antibodies by sequence similarity.

**Figure 1.**
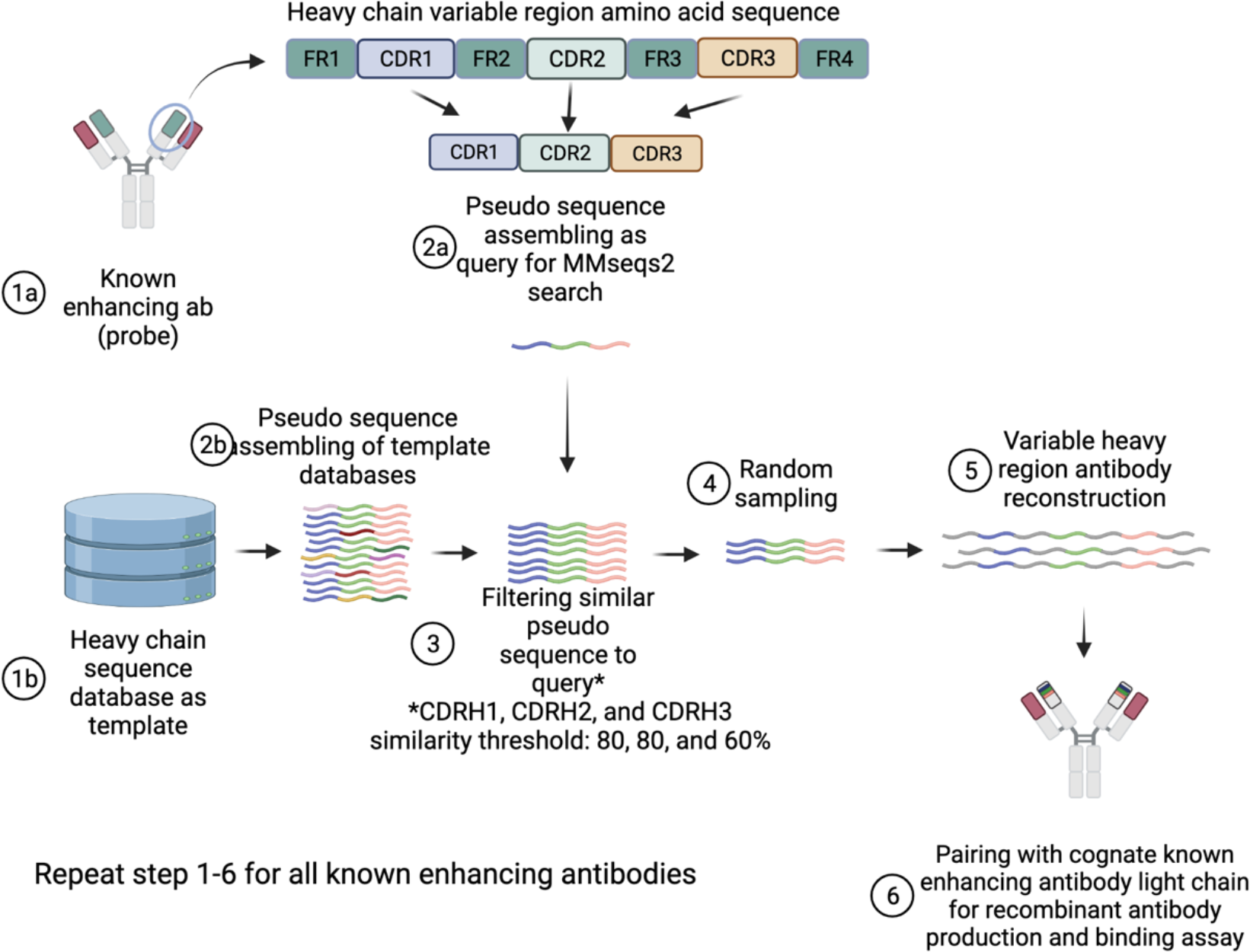
Schematic illustration of bioinformatics pipeline for finding functionally similar antibodies.

## Results

### Diverse antibodies target a common infection-enhancing epitopes on spike protein NTD

Despite targeting overlapping epitopes, the 11 previously reported enhancing antibodies have emerged from different germline genes and possess highly diverse CDRH3 amino acid sequences (Table S1 and Figure S1). Each sequence was expressed as a human IgG1 monoclonal antibody using a mammalian expression system and confirmed to recognize the wildtype spike protein, WT NTD but not to an NTD mutant with known epitope residues substituted with Alanine (W64A, H66A, V213A, and R214A) or to the WT RBD (Figure S2A-D). Binding to the Delta variant of SARS-CoV-2 Spike protein was also confirmed but the known enhancing antibodies lost their binding to the Omicron variant, which has extensive NTD mutations (Figure S2E-F). Moreover, all monoclonal antibodies facilitated ACE2 binding to WT and Delta Spike protein, but not to the Omicron variant (Figure S3).

### Encoding healthy and COVID-19 antibody repertoires for CDR similarity search

A total of 10 studies of healthy antibody repertoires, 15 studies of COVID-19 repertoires, and 3 studies of BNT162b2 vaccinated donor repertoires were collected. The healthy donor RNA sequencing data consisted of 297 donors (Table S2), COVID-19 data consisted of 213 patients (Table S3), and BNT162b2 vaccinated data consisted of 29 donors (Table S4). The data were processed to facilitate an efficient search by CDR similarity. A pseudo-sequence of concatenated CDRH1-3 amino acids was encoded as a MMseqs2 database (Steinegger & Soding, 2017). The resulting databases contained 55,401,329 healthy unvaccinated, 391,201 healthy BNT162b2 vaccinated, and 8,490,653 COVID-19 data entries, each linked to a complete variable region amino acid sequence.

### Sequences similar to enhancing antibodies found in healthy and COVID-19 repertoires

One of our motivations was to understand the relationship between CDRH3 sequence identity and shared epitope. We therefore searched for BCR sequences with rather loose criteria: where both CDRH1 and CDRH2 identity were at least 80% and CDRH3 was at least 60%. This search resulted in 7321 hits from healthy donors, 4679 from COVID-19 patients, and 113 from BNT162b2 vaccinated donors (Figure 2A). The distributions of CDR sequence identities among hits were similar for healthy unvaccinated, healthy vaccinated, and COVID-19 donors (Figure 2B). The CDRH3 sequences in COVID-19 hits were slightly more similar than those in healthy unvaccinated or healthy vaccinated donors. However, there were no heavy chain sequences that had the exact same amino acid sequence to a known enhancing antibody. B cells are known to be expanded and to acquire mutations upon antigen exposure to increase their affinity to antigens (Jacob et al., 1991). Indeed, among the hits, antigen exposed donors (COVID-19 and healthy vaccinated donors) had more expanded clones than healthy unvaccinated donors (Figure 2C). Sequences that were similar to COV2-2210, COV2-2582, and DH1055 dominated the COVID-19 and healthy unvaccinated donor hits (Figure 2D). In the healthy vaccinated donors, sequences that were similar to DH1054 were more dominant than sequences similar to COV2-2582. Sequences that were similar to DH1052, which has the longest CDRH3, were very infrequent in all the datasets.

**Figure 2.**
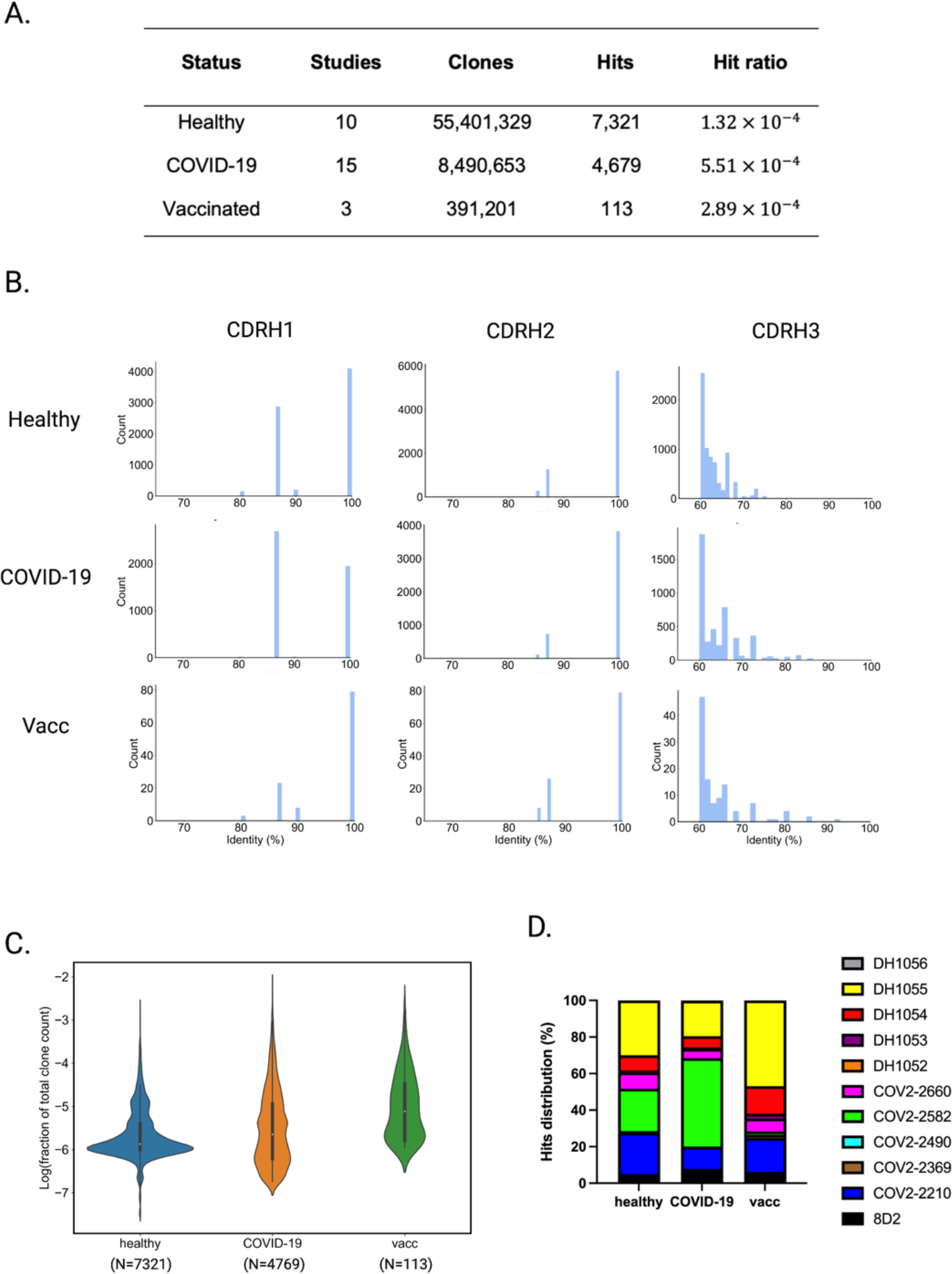
Finding enhancing antibodies from BCR sequencing data. (A) Enhancing antibodies hits from BCR sequencing data of healthy unvaccinated, COVID-19, and healthy vaccinated donors. (B) Hits CDRH1-3 identity distribution. (C) Fraction of total clone count of hits. The violin plot shows the distribution of fraction of total clone count with the higher fraction represent clonal expansion. (D) Distribution of enhancing antibodies hits based on the similarity to known enhancing antibodies.

### NTD-binding antibodies found in healthy donors and COVID-19 repertoire search hits

We next performed random sampling from healthy unvaccinated and COVID-19 search hits at a confidence level of 95% and a margin of error of 10%. Due to the low number of original sequences, the healthy vaccinated donors were excluded from the validation step. This resulted in selection of 96 non-redundant heavy chains from healthy unvaccinated donors and 94 from COVID-19 patients. The sampling qualitatively reproduced the original CDR similarity distribution (Figure 3A). The sequences that were similar to known infection-enhancing antibodies were then screened experimentally to observe whether they bound to the enhancing epitope or not. We observed that 17 out of 94 antibodies from COVID-19 donors (Figure 3B) and 2 out of 96 from healthy unvaccinated donors (Figure 3C) bound to S NTD, but not to the NTD mutant or the RBD (Figure S4 and S5). The fraction of antibodies from COVID-19 donors that bound to the S NTD or to the enhancing epitope was significantly higher than that in healthy unvaccinated donors (Chi-square test p-value < 0.01) (Figure 3D). Some antibodies exhibited higher binding affinity to the Spike protein from the Delta variant, but most lost their ability to bind to the Omicron variant (Figure S4D-E and S5D-E).

**Figure 3.**
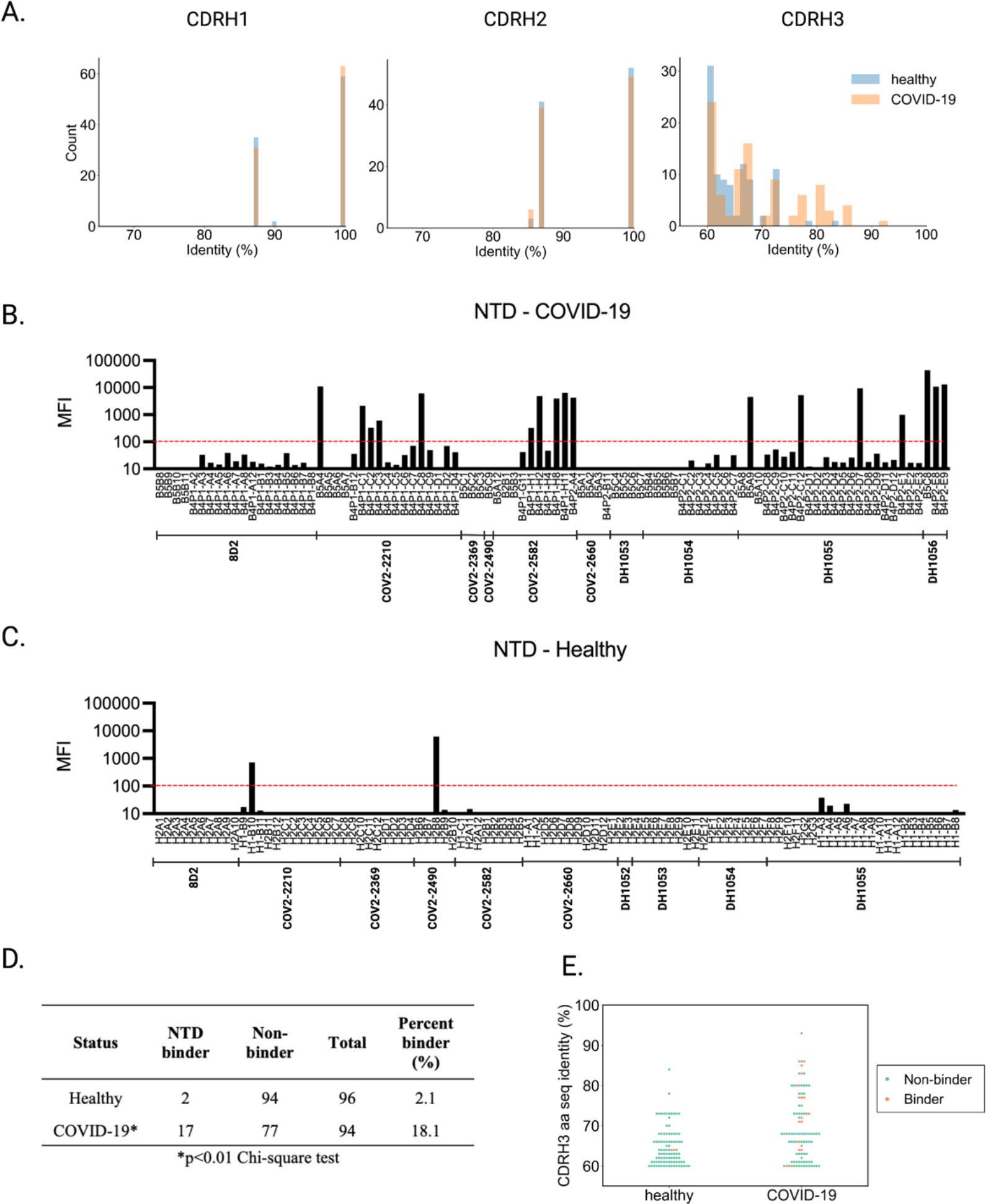
Sampling and testing the binding of hits to NTD. (A) CDRH1-3 distribution of sampled heavy chains. (B) NTD binding of produced antibodies from COVID-19 patients. Bars above the red dashed line are considered to be NTD binders. (C) NTD binding of produced antibodies from healthy unvaccinated donors. (D) NTD binders found from sampled antibodies and binder true positive rate for healthy unvaccinated and COVID-19. (E) CDRH3 distribution of NTD binders and non-binders.

Based on the true binders obtained from COVID-19 donors, the probability to find true binders among hits was approximately 30% for CDRH3 amino acid sequence identities above 70% but dropped to below 10% with CDRH3 sequence identities less than 70% (Figure 3E). In general, the binders and non-binders could not be separated by a single sequence identity cutoff. Additional sequence or structural data may allow us to differentiate binders from non-binders more accurately. The current results support the use of loose sequence identity thresholds when searching large repertoire data.

### A subset of antibodies that bind enhancing epitope enhance ACE2 binding

The previously known enhancing antibodies were able to increase ACE2 binding to Spike protein; therefore, we next assessed ACE2 binding enhancement using soluble ACE2 and Spike protein expressed on Expi293F cells (Liu et al., 2021). Antibodies that bound the enhancing epitope were purified and tested at the same concentration to confirm whether they increased the binding of ACE2 to the Spike protein. Among 19 antibodies tested, we found that 6 were potent ACE2 binding enhancers to Spike WT protein (Figure 4A). The ACE2 binding enhancement was increased when we tested these antibodies against Spike Delta variant, but they lost their ability when tested against Omicron variant (Figure 4B-C).

**Figure 4.**
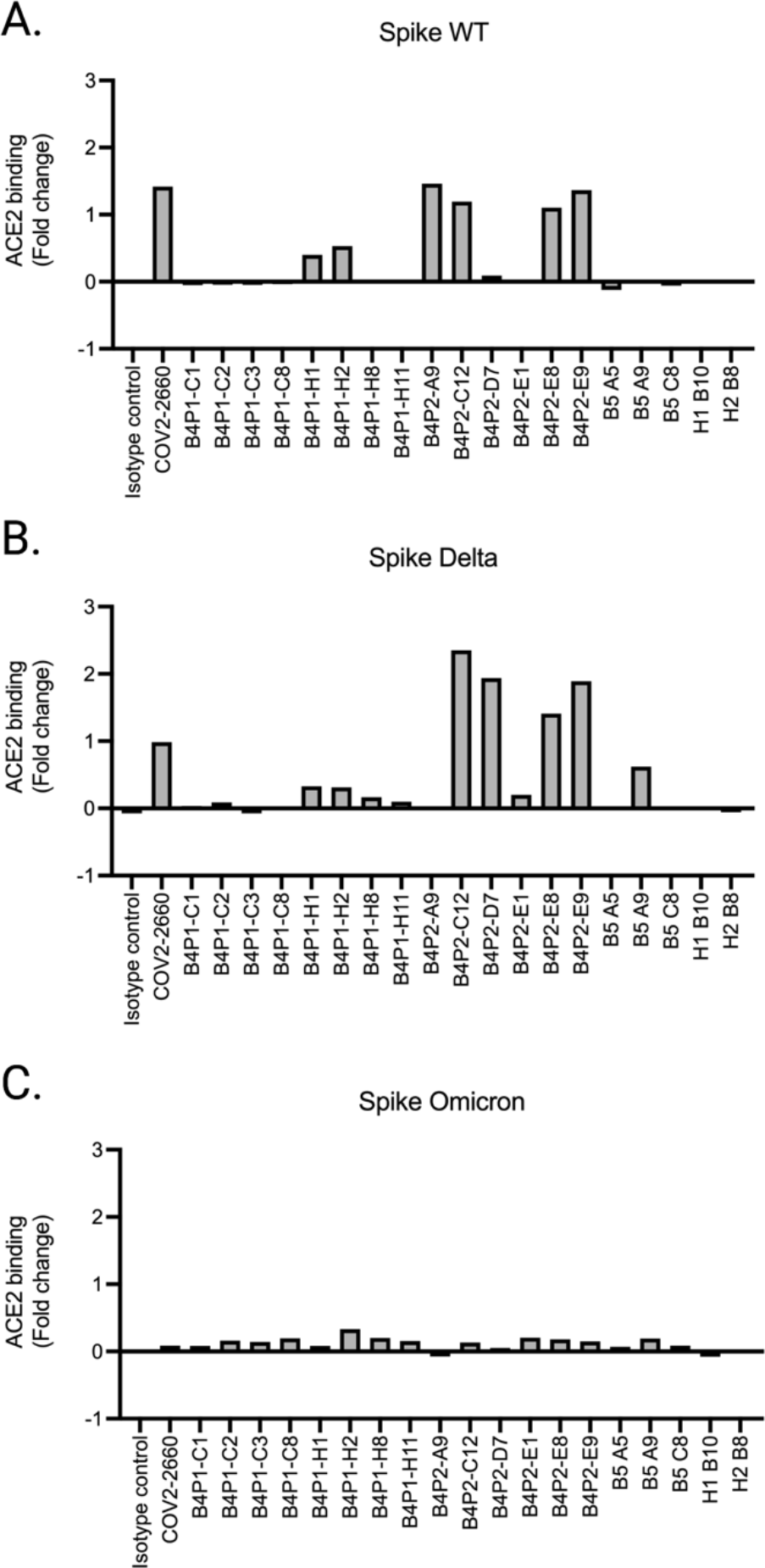
Enhancement of ACE2 binding to Spike protein in the present of antibodies. ACE2 binding to Spike WT (A), Delta (B), Omicron variant (C) enhancement are observed in the present of sampled antibodies or isotype control (hIgG1).

## Discussion

The observation that a subset of antibodies produced in diverse donors target an overlapping epitope, and that a further subset of the antibodies that binding to this epitope enhance ACE2 binding, raises several questions. We first sought to understand the distribution of these antibodies in both antigen-exposed and unexposed donors. Based on CDR sequence-similarity alone, we found highly similar distributions in healthy unvaccinated, healthy vaccinated and COVID-19 donors (Figure 2B). However, the clone sizes of these hits were qualitatively different in antigen-exposed and unexposed donors (Figure 2C). Furthermore, based on the experimental results, the frequency of binding to the enhancing epitope was significantly higher for the COVID-19 derived group than the healthy-derived group. From the NTD binding data, we could estimate the frequency of enhancing antibodies within COVID-19 and healthy unvaccinated donors to be less than 100 and 3 per million clones, respectively. Although the proposed enhancing mechanism requires a more detailed study, it may well apply to other coronaviruses that use ACE2 as a host receptor. Given the large reservoir of potential antibodies in the healthy population, this may represent a modest concern for future vaccine design.

Ease of access to RNA sequencing technologies, as well as reduction of cost has resulted in a rapid increase in publicly available BCR repertoire sequence data (Marks & Deane, 2020). The approach taken here to search this data is general and will likely aid in the discovery of not only enhancing, but also neutralizing antibodies, autoantibodies, or even T cell receptors. As more data on binders and non-binders accumulates, more sophisticated metrics of similarity can be tested. The methods used here are available as an open-source project with a freely accessible web server (www.sysimm.org/interclone/).

## Acknowledgments

The authors would like to thank RIMD and IFReC Core Experimental Facility for the support in conducting experiments; all Standley Lab member for constructive discussion and comments on the manuscript. This work was supported by Japan Agency for Medical Research and Development (AMED), Platform Project for Supporting Drug Discovery and Life Science Research (Basis for Supporting Innovative Drug Discovery and Life Science Research) under JP21am0101108.

## Author contributions

H.S.I. performed searching of new enhancing antibodies, binding assay, ACE2 binding enhancement assay, and data analysis. H.S.I. and D.S.S. performed antibodies expression. Z.X. and J.W. developed the backend databases. Z.X., J.W., S.L. and D.M.S. did the bioinformatics pipeline development. H.S.I., D.M.S., D.K.N., Y.H., H.A., M.O. conceptualized and designed the experiments. All authors wrote, reviewed, and edited the manuscript. D.M.S. supervised the overall project.

## Method details

### BCR repertoire data mining and processing

Datasets were retrieved from publicly available repository such as sequence read archive (SRA) and European nucleotide archive (ENA). Collected study detail can be seen in the Table S2-S4 (Bernardes et al., 2020; Galson et al., 2020; Ghraichy et al., 2020; Gidoni et al., 2019; Goel, Apostolidis, et al., 2021; Goel, Painter, et al., 2021; Kim et al., 2021; Kuri-Cervantes et al., 2020; Meng et al., 2017; Montague et al., 2021; Mor et al., 2021; Nielsen et al., 2020; Niu et al., 2020; Roskin et al., 2020; Schmitz et al., 2021; Schultheiss et al., 2020; Setliff et al., 2018; Sokal et al., 2021; Soto et al., 2019; Turner et al., 2021; Turner et al., 2020; Wen et al., 2020; Woodruff et al., 2020; Zhang et al., 2020; Zhou et al., 2021). Raw BCR repertoire sequencing data was formatted into AIRR-formatted files and then CDRs was assigned using ANARCI (Dunbar & Deane, 2016). CDRH1, CDRH3, and CDRH3 that have been assigned then concatenated into single pseudo-sequences for later encoding into MMSeqs2 database format (Steinegger & Soding, 2017). Query sequences (11 enhancing antibodies) were also processed similarly. For each query sequence, database was searched using the minimum sequence identity cutoff. For each hit, pseudo-sequence was separated into CDRH1, CDRH2, and CDRH3 then sequence identity for each CDR was evaluated (CDRH1, CDRH2, and CDRH3 cutoff 80, 80, and 60 %).

### Cell lines

Expi293F cells (Thermo) were maintained in Expi293 expression medium (Gibco) supplemented with 100x dilution of 10,000 U/mL penicillin/streptomycin (Gibco) at 37°C incubators under 8% CO2 and shaking at 125 rpm.

### Production of infection enhancing antibodies from COVID-19 patients and healthy donors sequence database

Recombinant antibodies were produced as previously described (Liu et al., 2021). Briefly, the variable regions of sampled heavy chains from the COVID-19 patients and healthy donors were prepared by dsDNA synthesis (IDT) and cloned into pCAGGS vectors containing sequences of human IgG1 constant region. The light chains from known enhancing antibodies were synthesized and cloned into the pCAGGS vector containing the human immunoglobulin kappa constant region. To produce recombinant antibodies, vectors containing heavy chain sequence and light chain sequence from known antibodies were co-transfected into Expi293F cells (Thermo) and the supernatant was collected for further assay.

### Antibody binding assay

Antibody binding to SARS-CoV-2 antigen was measured as previously described (Liu et al., 2021) with modification of antigen display cells from HEK293T to Expi293F (Thermo). The pME18S plasmid expressing Spike protein C-terminal retention signal deletion from Wuhan (WT), Delta, and Omicron variant, Flag-NTD-PILR-TM, Flag-RBD-PILR-TM, and Flag-NTD (W64A, H66A, V213A, and R214A)-PILR-TM, were co-transfected with pMx plasmid expressing GFP as the marker to the Expi293F cells (Thermo). The transfectant cells were incubated with supernatant containing expressed antibodies for 30 minutes then followed by incubation with APC-anti-human IgG (H+L) antibody (Jackson ImmunoResearch, USA). Bound antibodies to the GFP-positive cells were then analyzed by flow cytometry (Attune NxT, Thermo).

### ACE2 binding enhancement assay

The SARS-CoV-2 S-NTD binders were expressed and purified using protein A spin column (Cosmo Bio) and concentration was measured using ELISA. ACE2 binding enhancement to Spike WT, Delta, and Omicron variants in the presence of enhancing antibodies was measured. Briefly, Expi293F cells (Thermo) that express either Spike WT, Delta, or Omicron variant were incubated by 1 μg/mL antibodies for 30 minutes. Followed by ACE2-biotin at 1 μg/mL (RnD Systems) incubation for 30 minutes and then SA-APC (Biolegend) incubation for 1 hour. The amount of ACE2 that binds to Spike protein was measured using flow cytometry (Attune NxT, Thermo). Fold change was calculated by comparing the amount of bound ACE2 in the presence of antibodies and in the absence of antibodies.

### Quantification and Statistical Analysis

Flow cytometry data were analyzed using FlowJo version 10.7 (BD Biosciences, USA). GraphPad Prism version 9 was used for binding assay graph generation. Matplotlib (v. 3.3.4) and Seaborn (v. 0.11.0) python packages were used to generate CDRs similarity and ACE2 enhancement graph and violin plot. Scipy (v. 1.7.3) was used to calculate Chi-square test.

## References

Akkaya, M., Kwak, K., & Pierce, S. K. (2020). B cell memory: building two walls of protection against pathogens. Nat Rev Immunol, 20(4), 229–238. https://doi.org/10.1038/s41577-019-0244-2

Arvin, A. M., Fink, K., Schmid, M. A., Cathcart, A., Spreafico, R., Havenar-Daughton, C., Lanzavecchia, A., Corti, D., & Virgin, H. W. (2020). A perspective on potential antibody-dependent enhancement of SARS-CoV-2. Nature, 584(7821), 353–363. https://doi.org/10.1038/s41586-020-2538-8

Bernardes, J. P., Mishra, N., Tran, F., Bahmer, T., Best, L., Blase, J. I., Bordoni, D., Franzenburg, J., Geisen, U., Josephs-Spaulding, J., Kohler, P., Kunstner, A., Rosati, E., Aschenbrenner, A. C., Bacher, P., Baran, N., Boysen, T., Brandt, B., Bruse, N., … Deutsche, C.-O. I. (2020). Longitudinal Multi-omics Analyses Identify Responses of Megakaryocytes, Erythroid Cells, and Plasmablasts as Hallmarks of Severe COVID-19. Immunity, 53(6), 1296–1314 e1299. https://doi.org/10.1016/j.immuni.2020.11.017

Bournazos, S., Gupta, A., & Ravetch, J. V. (2020). The role of IgG Fc receptors in antibody-dependent enhancement. Nat Rev Immunol, 20(10), 633–643. https://doi.org/10.1038/s41577-020-00410-0

Cyster, J. G., & Allen, C. D. C. (2019). B Cell Responses: Cell Interaction Dynamics and Decisions. Cell, 177(3), 524–540. https://doi.org/10.1016/j.cell.2019.03.016

Dorner, T., & Radbruch, A. (2007). Antibodies and B cell memory in viral immunity. Immunity, 27(3), 384–392. https://doi.org/10.1016/j.immuni.2007.09.002

Dunbar, J., & Deane, C. M. (2016). ANARCI: antigen receptor numbering and receptor classification. Bioinformatics, 32(2), 298–300. https://doi.org/10.1093/bioinformatics/btv552

Galson, J. D., Schaetzle, S., Bashford-Rogers, R. J. M., Raybould, M. I. J., Kovaltsuk, A., Kilpatrick, G. J., Minter, R., Finch, D. K., Dias, J., James, L. K., Thomas, G., Lee, W. J., Betley, J., Cavlan, O., Leech, A., Deane, C. M., Seoane, J., Caldas, C., Pennington, D. J., … Osbourn, J. (2020). Deep Sequencing of B Cell Receptor Repertoires From COVID-19 Patients Reveals Strong Convergent Immune Signatures. Front Immunol, 11, 605170. https://doi.org/10.3389/fimmu.2020.605170

Ghraichy, M., Galson, J. D., Kovaltsuk, A., von Niederhausern, V., Pachlopnik Schmid, J., Recher, M., Jauch, A. J., Miho, E., Kelly, D. F., Deane, C. M., & Truck, J. (2020). Maturation of the Human Immunoglobulin Heavy Chain Repertoire With Age. Front Immunol, 11, 1734. https://doi.org/10.3389/fimmu.2020.01734

Gidoni, M., Snir, O., Peres, A., Polak, P., Lindeman, I., Mikocziova, I., Sarna, V. K., Lundin, K. E. A., Clouser, C., Vigneault, F., Collins, A. M., Sollid, L. M., & Yaari, G. (2019). Mosaic deletion patterns of the human antibody heavy chain gene locus shown by Bayesian haplotyping. Nat Commun, 10(1), 628. https://doi.org/10.1038/s41467-019-08489-3

Goel, R. R., Apostolidis, S. A., Painter, M. M., Mathew, D., Pattekar, A., Kuthuru, O., Gouma, S., Hicks, P., Meng, W., Rosenfeld, A. M., Dysinger, S., Lundgreen, K. A., Kuri-Cervantes, L., Adamski, S., Hicks, A., Korte, S., Oldridge, D. A., Baxter, A. E., Giles, J. R., … Wherry, E. J. (2021). Distinct antibody and memory B cell responses in SARS-CoV-2 naive and recovered individuals following mRNA vaccination. Sci Immunol, 6(58). https://doi.org/10.1126/sciimmunol.abi6950

Goel, R. R., Painter, M. M., Apostolidis, S. A., Mathew, D., Meng, W., Rosenfeld, A. M., Lundgreen, K. A., Reynaldi, A., Khoury, D. S., Pattekar, A., Gouma, S., Kuri-Cervantes, L., Hicks, P., Dysinger, S., Hicks, A., Sharma, H., Herring, S., Korte, S., Baxter, A. E., … Wherry, E. J. (2021). mRNA vaccines induce durable immune memory to SARS-CoV-2 and variants of concern. Science, 374(6572), abm0829. https://doi.org/10.1126/science.abm0829

Guzman, M. G., Alvarez, M., & Halstead, S. B. (2013). Secondary infection as a risk factor for dengue hemorrhagic fever/dengue shock syndrome: an historical perspective and role of antibody-dependent enhancement of infection. Arch Virol, 158(7), 1445–1459. https://doi.org/10.1007/s00705-013-1645-3

Harwood, N. E., & Batista, F. D. (2010). Early events in B cell activation. Annu Rev Immunol, 28, 185–210. https://doi.org/10.1146/annurev-immunol-030409-101216

Haynes, B. F., Corey, L., Fernandes, P., Gilbert, P. B., Hotez, P. J., Rao, S., Santos, M. R., Schuitemaker, H., Watson, M., & Arvin, A. (2020). Prospects for a safe COVID-19 vaccine. Sci Transl Med, 12(568). https://doi.org/10.1126/scitranslmed.abe0948

Jacob, J., Kelsoe, G., Rajewsky, K., & Weiss, U. (1991). Intraclonal generation of antibody mutants in germinal centres. Nature, 354(6352), 389–392. https://doi.org/10.1038/354389a0

Kapikian, A. Z., Mitchell, R. H., Chanock, R. M., Shvedoff, R. A., & Stewart, C. E. (1969). An epidemiologic study of altered clinical reactivity to respiratory syncytial (RS) virus infection in children previously vaccinated with an inactivated RS virus vaccine. Am J Epidemiol, 89(4), 405–421. https://doi.org/10.1093/oxfordjournals.aje.a120954

Kim, H. W., Canchola, J. G., Brandt, C. D., Pyles, G., Chanock, R. M., Jensen, K., & Parrott, R. H. (1969). Respiratory syncytial virus disease in infants despite prior administration of antigenic inactivated vaccine. Am J Epidemiol, 89(4), 422–434. https://doi.org/10.1093/oxfordjournals.aje.a120955

Kim, S. I., Noh, J., Kim, S., Choi, Y., Yoo, D. K., Lee, Y., Lee, H., Jung, J., Kang, C. K., Song, K. H., Choe, P. G., Kim, H. B., Kim, E. S., Kim, N. J., Seong, M. W., Park, W. B., Oh, M. D., Kwon, S., & Chung, J. (2021). Stereotypic neutralizing VH antibodies against SARS-CoV-2 spike protein receptor binding domain in patients with COVID-19 and healthy individuals. Sci Transl Med, 13(578). https://doi.org/10.1126/scitranslmed.abd6990

Kuri-Cervantes, L., Pampena, M. B., Meng, W., Rosenfeld, A. M., Ittner, C. A. G., Weisman, A. R., Agyekum, R. S., Mathew, D., Baxter, A. E., Vella, L. A., Kuthuru, O., Apostolidis, S. A., Bershaw, L., Dougherty, J., Greenplate, A. R., Pattekar, A., Kim, J., Han, N., Gouma, S., … Betts, M. R. (2020). Comprehensive mapping of immune perturbations associated with severe COVID-19. Sci Immunol, 5(49). https://doi.org/10.1126/sciimmunol.abd7114

Kurosaki, T., Kometani, K., & Ise, W. (2015). Memory B cells. Nat Rev Immunol, 15(3), 149–159. https://doi.org/10.1038/nri3802

Li, D., Edwards, R. J., Manne, K., Martinez, D. R., Schafer, A., Alam, S. M., Wiehe, K., Lu, X., Parks, R., Sutherland, L. L., Oguin, T. H., 3rd, McDanal, C., Perez, L. G., Mansouri, K., Gobeil, S. M. C., Janowska, K., Stalls, V., Kopp, M., Cai, F., … Saunders, K. O. (2021). In vitro and in vivo functions of SARS-CoV-2 infection-enhancing and neutralizing antibodies. Cell, 184(16), 4203–4219 e4232. https://doi.org/10.1016/j.cell.2021.06.021

Liu, Y., Soh, W. T., Kishikawa, J. I., Hirose, M., Nakayama, E. E., Li, S., Sasai, M., Suzuki, T., Tada, A., Arakawa, A., Matsuoka, S., Akamatsu, K., Matsuda, M., Ono, C., Torii, S., Kishida, K., Jin, H., Nakai, W., Arase, N., … Arase, H. (2021). An infectivity-enhancing site on the SARS-CoV-2 spike protein targeted by antibodies. Cell, 184(13), 3452–3466 e3418. https://doi.org/10.1016/j.cell.2021.05.032

Marks, C., & Deane, C. M. (2020). How repertoire data are changing antibody science. J Biol Chem, 295(29), 9823–9837. https://doi.org/10.1074/jbc.REV120.010181

Meng, W., Zhang, B., Schwartz, G. W., Rosenfeld, A. M., Ren, D., Thome, J. J. C., Carpenter, D. J., Matsuoka, N., Lerner, H., Friedman, A. L., Granot, T., Farber, D. L., Shlomchik, M. J., Hershberg, U., & Luning Prak, E. T. (2017). An atlas of B-cell clonal distribution in the human body. Nat Biotechnol, 35(9), 879–884. https://doi.org/10.1038/nbt.3942

Montague, Z., Lv, H., Otwinowski, J., DeWitt, W. S., Isacchini, G., Yip, G. K., Ng, W. W., Tsang, O. T., Yuan, M., Liu, H., Wilson, I. A., Peiris, J. S. M., Wu, N. C., Nourmohammad, A., & Mok, C. K. P. (2021). Dynamics of B cell repertoires and emergence of cross-reactive responses in patients with different severities of COVID-19. Cell Rep, 35(8), 109173. https://doi.org/10.1016/j.celrep.2021.109173

Mor, M., Werbner, M., Alter, J., Safra, M., Chomsky, E., Lee, J. C., Hada-Neeman, S., Polonsky, K., Nowell, C. J., Clark, A. E., Roitburd-Berman, A., Ben-Shalom, N., Navon, M., Rafael, D., Sharim, H., Kiner, E., Griffis, E. R., Gershoni, J. M., Kobiler, O., … Freund, N. T. (2021). Multi-clonal SARS-CoV-2 neutralization by antibodies isolated from severe COVID-19 convalescent donors. PLoS Pathog, 17(2), e1009165. https://doi.org/10.1371/journal.ppat.1009165

Morales-Nunez, J. J., Munoz-Valle, J. F., Torres-Hernandez, P. C., & Hernandez-Bello, J. (2021). Overview of Neutralizing Antibodies and Their Potential in COVID-19. Vaccines (Basel), 9(12). https://doi.org/10.3390/vaccines9121376

Nielsen, S. C. A., Yang, F., Jackson, K. J. L., Hoh, R. A., Roltgen, K., Jean, G. H., Stevens, B. A., Lee, J. Y., Rustagi, A., Rogers, A. J., Powell, A. E., Hunter, M., Najeeb, J., Otrelo-Cardoso, A. R., Yost, K. E., Daniel, B., Nadeau, K. C., Chang, H. Y., Satpathy, A. T., … Boyd, S. D. (2020). Human B Cell Clonal Expansion and Convergent Antibody Responses to SARS-CoV-2. Cell Host Microbe, 28(4), 516–525 e515. https://doi.org/10.1016/j.chom.2020.09.002

Niu, X., Li, S., Li, P., Pan, W., Wang, Q., Feng, Y., Mo, X., Yan, Q., Ye, X., Luo, J., Qu, L., Weber, D., Byrne-Steele, M. L., Wang, Z., Yu, F., Li, F., Myers, R. M., Lotze, M. T., Zhong, N., … Chen, L. (2020). Longitudinal Analysis of T and B Cell Receptor Repertoire Transcripts Reveal Dynamic Immune Response in COVID-19 Patients. Front Immunol, 11, 582010. https://doi.org/10.3389/fimmu.2020.582010

Phan, T. G., & Tangye, S. G. (2017). Memory B cells: total recall. Curr Opin Immunol, 45, 132–140. https://doi.org/10.1016/j.coi.2017.03.005

Polack, F. P., Hoffman, S. J., Crujeiras, G., & Griffin, D. E. (2003). A role for nonprotective complement-fixing antibodies with low avidity for measles virus in atypical measles. Nat Med, 9(9), 1209–1213. https://doi.org/10.1038/nm918

Roskin, K. M., Jackson, K. J. L., Lee, J. Y., Hoh, R. A., Joshi, S. A., Hwang, K. K., Bonsignori, M., Pedroza-Pacheco, I., Liao, H. X., Moody, M. A., Fire, A. Z., Borrow, P., Haynes, B. F., & Boyd, S. D. (2020). Aberrant B cell repertoire selection associated with HIV neutralizing antibody breadth. Nat Immunol, 21(2), 199–209. https://doi.org/10.1038/s41590-019-0581-0

Schmitz, A. J., Turner, J. S., Liu, Z., Zhou, J. Q., Aziati, I. D., Chen, R. E., Joshi, A., Bricker, T. L., Darling, T. L., Adelsberg, D. C., Altomare, C. G., Alsoussi, W. B., Case, J. B., VanBlargan, L. A., Lei, T., Thapa, M., Amanat, F., Jeevan, T., Fabrizio, T., … Ellebedy, A. H. (2021). A vaccine-induced public antibody protects against SARS-CoV-2 and emerging variants. Immunity, 54(9), 2159–2166 e2156. https://doi.org/10.1016/j.immuni.2021.08.013

Schultheiss, C., Paschold, L., Simnica, D., Mohme, M., Willscher, E., von Wenserski, L., Scholz, R., Wieters, I., Dahlke, C., Tolosa, E., Sedding, D. G., Ciesek, S., Addo, M., & Binder, M. (2020). Next-Generation Sequencing of T and B Cell Receptor Repertoires from COVID-19 Patients Showed Signatures Associated with Severity of Disease. Immunity, 53(2), 442–455 e444. https://doi.org/10.1016/j.immuni.2020.06.024

Setliff, I., McDonnell, W. J., Raju, N., Bombardi, R. G., Murji, A. A., Scheepers, C., Ziki, R., Mynhardt, C., Shepherd, B. E., Mamchak, A. A., Garrett, N., Karim, S. A., Mallal, S. A., Crowe, J. E., Jr., Morris, L., & Georgiev, I. S. (2018). Multi-Donor Longitudinal Antibody Repertoire Sequencing Reveals the Existence of Public Antibody Clonotypes in HIV-1 Infection. Cell Host Microbe, 23(6), 845–854 e846. https://doi.org/10.1016/j.chom.2018.05.001

Simmons, C. P., Farrar, J. J., Nguyen v, V., & Wills, B. (2012). Dengue. N Engl J Med, 366(15), 1423–1432. https://doi.org/10.1056/NEJMra1110265

Sokal, A., Chappert, P., Barba-Spaeth, G., Roeser, A., Fourati, S., Azzaoui, I., Vandenberghe, A., Fernandez, I., Meola, A., Bouvier-Alias, M., Crickx, E., Beldi-Ferchiou, A., Hue, S., Languille, L., Michel, M., Baloul, S., Noizat-Pirenne, F., Luka, M., Megret, J., … Mahevas, M. (2021). Maturation and persistence of the anti-SARS-CoV-2 memory B cell response. Cell, 184(5), 1201–1213 e1214. https://doi.org/10.1016/j.cell.2021.01.050

Soto, C., Bombardi, R. G., Branchizio, A., Kose, N., Matta, P., Sevy, A. M., Sinkovits, R. S., Gilchuk, P., Finn, J. A., & Crowe, J. E., Jr. (2019). High frequency of shared clonotypes in human B cell receptor repertoires. Nature, 566(7744), 398–402. https://doi.org/10.1038/s41586-019-0934-8

Steinegger, M., & Soding, J. (2017). MMseqs2 enables sensitive protein sequence searching for the analysis of massive data sets. Nat Biotechnol, 35(11), 1026–1028. https://doi.org/10.1038/nbt.3988

Turner, J. S., O’Halloran, J. A., Kalaidina, E., Kim, W., Schmitz, A. J., Zhou, J. Q., Lei, T., Thapa, M., Chen, R. E., Case, J. B., Amanat, F., Rauseo, A. M., Haile, A., Xie, X., Klebert, M. K., Suessen, T., Middleton, W. D., Shi, P. Y., Krammer, F., … Ellebedy, A. H. (2021). SARS-CoV-2 mRNA vaccines induce persistent human germinal centre responses. Nature, 596(7870), 109–113. https://doi.org/10.1038/s41586-021-03738-2

Turner, J. S., Zhou, J. Q., Han, J., Schmitz, A. J., Rizk, A. A., Alsoussi, W. B., Lei, T., Amor, M., McIntire, K. M., Meade, P., Strohmeier, S., Brent, R. I., Richey, S. T., Haile, A., Yang, Y. R., Klebert, M. K., Suessen, T., Teefey, S., Presti, R. M., … Ellebedy, A. H. (2020). Human germinal centres engage memory and naive B cells after influenza vaccination. Nature, 586(7827), 127–132. https://doi.org/10.1038/s41586-020-2711-0

Victora, G. D., & Nussenzweig, M. C. (2012). Germinal centers. Annu Rev Immunol, 30, 429–457. https://doi.org/10.1146/annurev-immunol-020711-075032

Wen, W., Su, W., Tang, H., Le, W., Zhang, X., Zheng, Y., Liu, X., Xie, L., Li, J., Ye, J., Dong, L., Cui, X., Miao, Y., Wang, D., Dong, J., Xiao, C., Chen, W., & Wang, H. (2020). Immune cell profiling of COVID-19 patients in the recovery stage by single-cell sequencing. Cell Discov, 6, 31. https://doi.org/10.1038/s41421-020-0168-9

Woodruff, M. C., Ramonell, R. P., Nguyen, D. C., Cashman, K. S., Saini, A. S., Haddad, N. S., Ley, A. M., Kyu, S., Howell, J. C., Ozturk, T., Lee, S., Suryadevara, N., Case, J. B., Bugrovsky, R., Chen, W., Estrada, J., Morrison-Porter, A., Derrico, A., Anam, F. A., … Sanz, I. (2020). Extrafollicular B cell responses correlate with neutralizing antibodies and morbidity in COVID-19. Nat Immunol, 21(12), 1506–1516. https://doi.org/10.1038/s41590-020-00814-z

Zhang, J. Y., Wang, X. M., Xing, X., Xu, Z., Zhang, C., Song, J. W., Fan, X., Xia, P., Fu, J. L., Wang, S. Y., Xu, R. N., Dai, X. P., Shi, L., Huang, L., Jiang, T. J., Shi, M., Zhang, Y., Zumla, A., Maeurer, M., … Wang, F. S. (2020). Single-cell landscape of immunological responses in patients with COVID-19. Nat Immunol, 21(9), 1107–1118. https://doi.org/10.1038/s41590-020-0762-x

Zhou, Y., Zhang, J., Wang, D., Wang, D., Guan, W., Qin, J., Xu, X., Fang, J., Fu, B., Zheng, X., Wang, D., Zhao, H., Chen, X., Tian, Z., Xu, X., Wang, G., & Wei, H. (2021). Profiling of the immune repertoire in COVID-19 patients with mild, severe, convalescent, or retesting-positive status. J Autoimmun, 118, 102596. https://doi.org/10.1016/j.jaut.2021.102596

