## Supplementary Information for "Landscape of infection enhancing antibodies in COVID-19 and healthy donors"

* Corresponding author

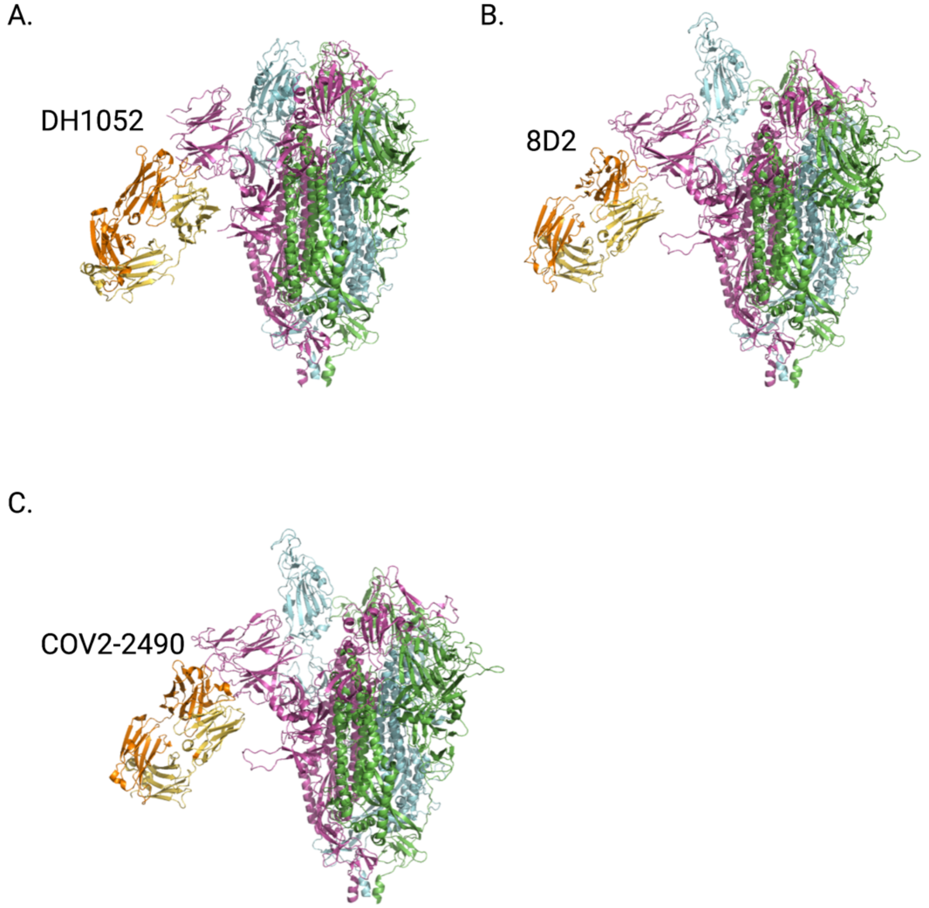


**Figure S1. Cryo-em protein structure representation of enhancing antibodies binding to Spike protein**

(A) Structure of DH1052 binds to Spike protein (PDB ID: 7LAB). (B) Structure of 8D2 binds to Spike protein (PDB ID: 7DZX). (C) Structure of COV2-2490 binds to Spike protein (PDB ID: 7DZY).

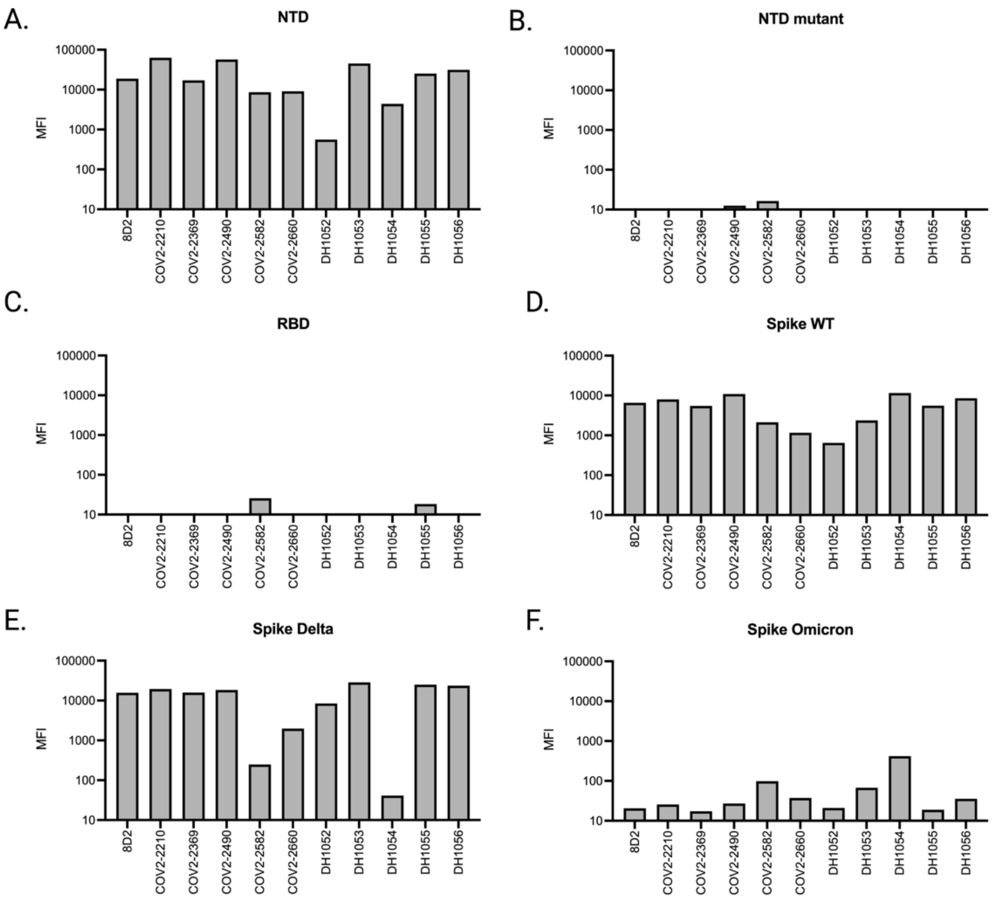


**Figure S2. Known enhancing antibody binding to various SARS-CoV-2 antigen**

Antibodies binding to NTD (A), NTD mutant (B), RBD (C), Spike WT (D), Spike Delta (E), and Spike Omicron (F) are represented as Mean Fluorescence Intensity (MFI).

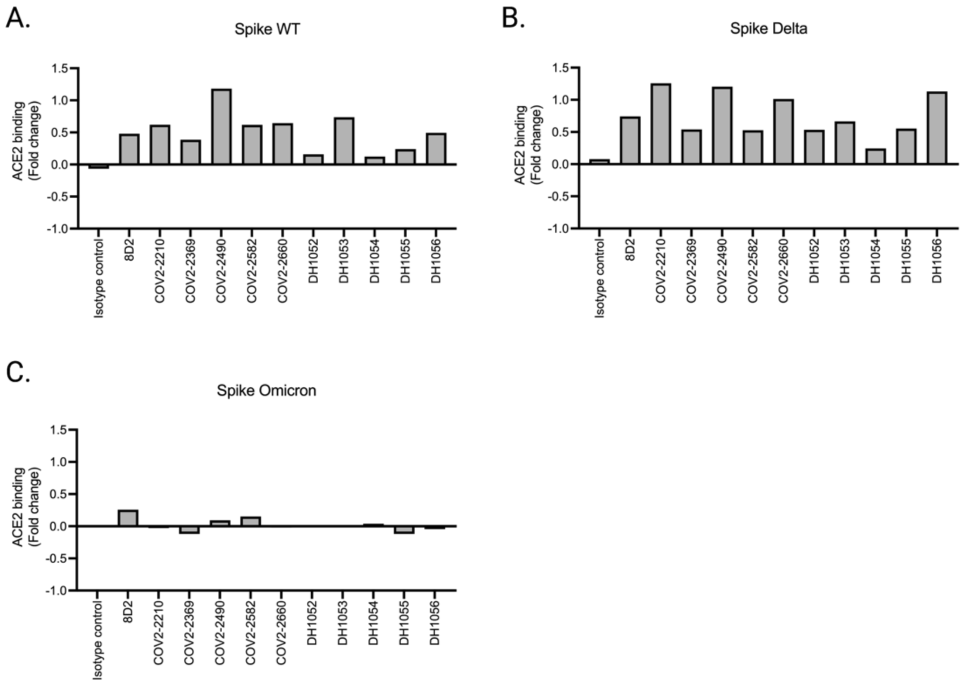


**Figure S3. Known enhancing antibodies enhance ACE2 binding to Spike protein**

Spike WT (A), Delta (B), and Omicron variant (C) were used as the antigen target for ACE2 binding. Fold change was calculated as ACE2 binding MFI in the presence of antibodies subtracted by ACE2 binding MFI without antibodies then divided by ACE2 binding MFI without MFI.

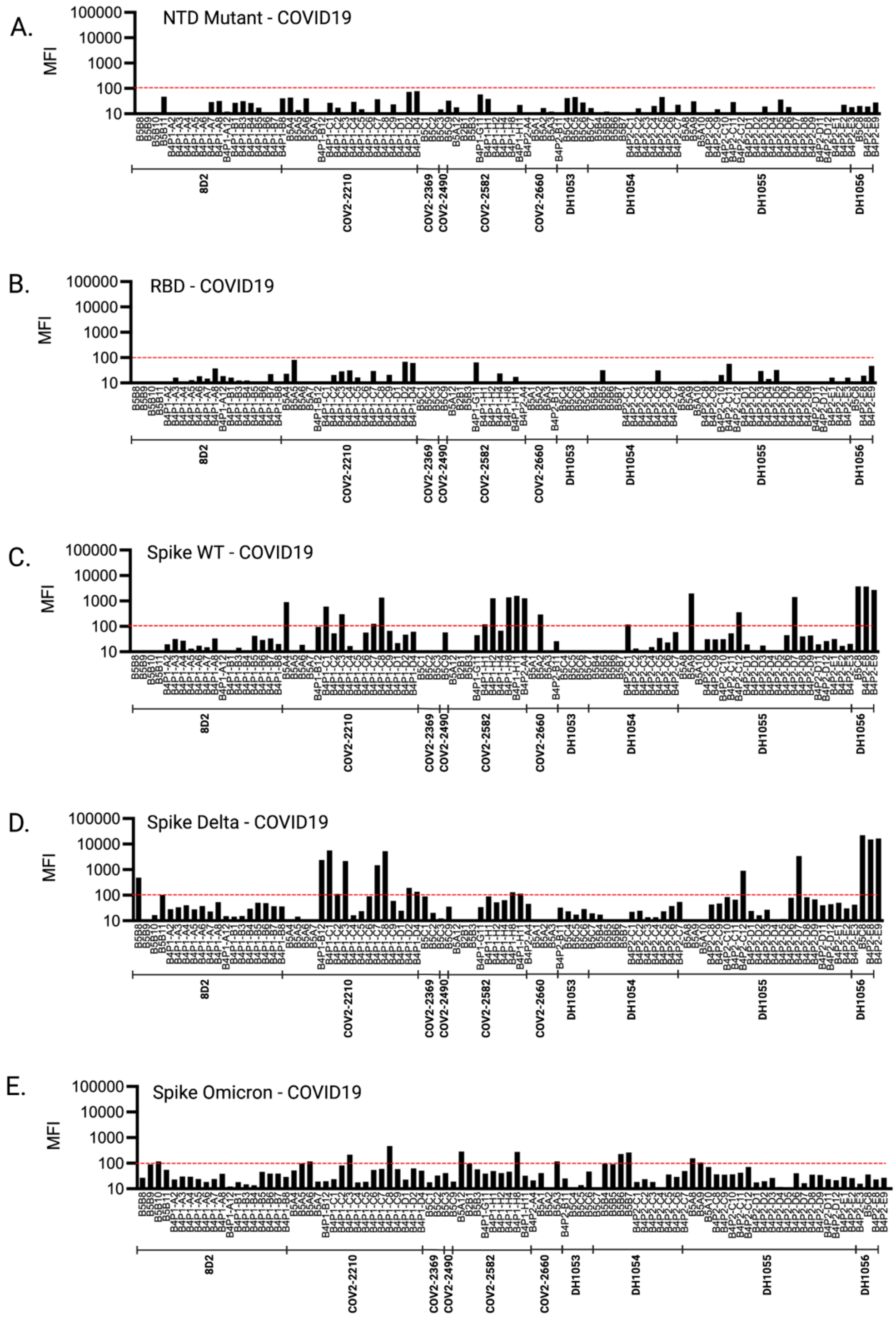


**Figure S4. Sampled antibodies binding to SARS-CoV-2 antigens from sampled COVID-19 donors' antibodies**

NTD mutant (A), RBD (B), Spike WT (C), Spike Delta (D), and Spike Omicron (E) and shown as MFI.

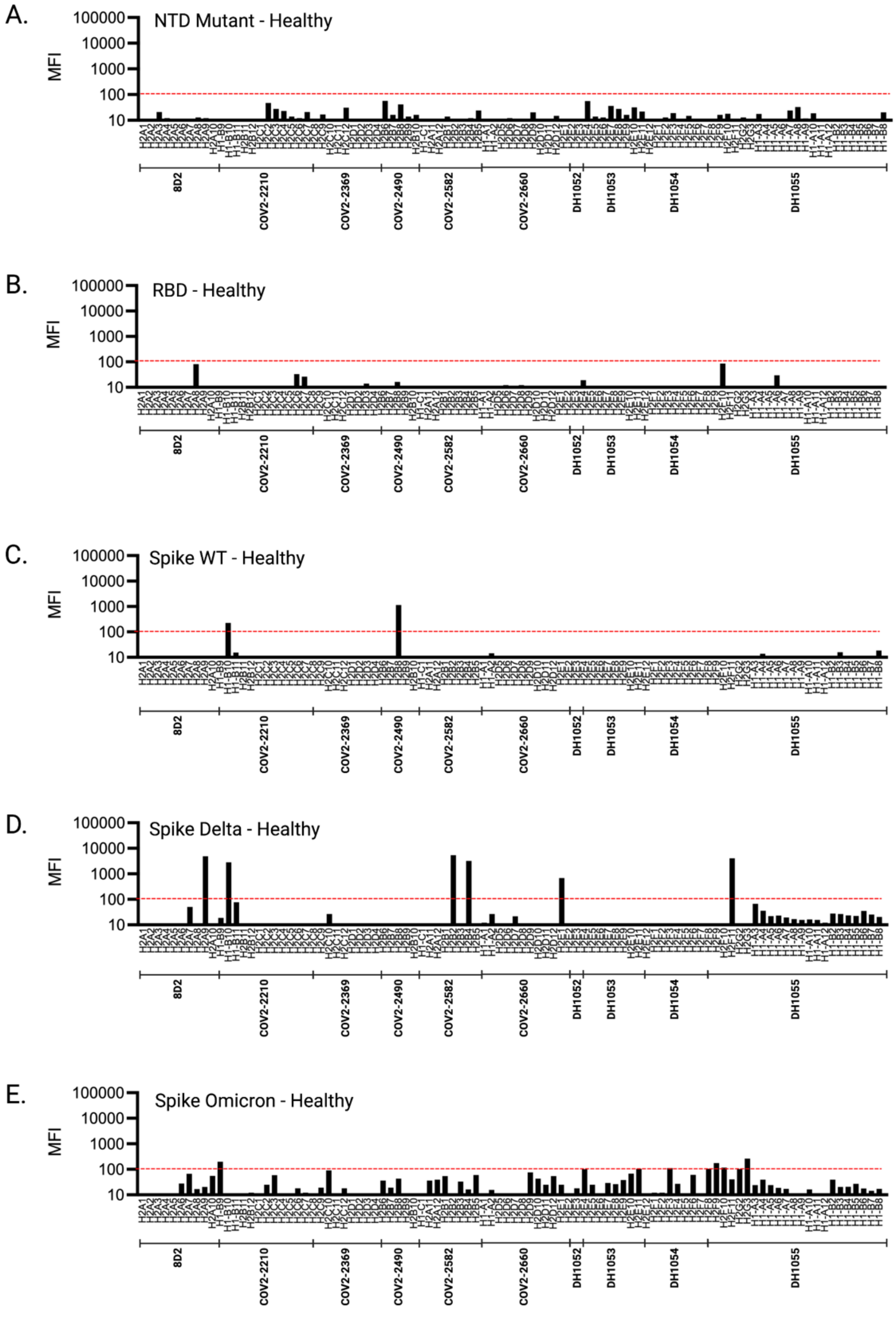


**Figure S4. Sampled antibodies binding to SARS-CoV-2 antigens from sampled healthy unvaccinated donors' antibodies**

NTD mutant (A), RBD (B), Spike WT (C), Spike Delta (D), and Spike Omicron (E) and shown as MFI.

**Table S1. Gene usage of known enhancing antibodies.**

| Name | $V_H$ gene | $J_H$ gene | CDRH3 aa | $V_L$ gene | $J_L$ gene | CDRL3 aa |
|---|---|---|---|---|---|---|
| 8D2 | IGHV3-7 | IGHJ3 | ARDWDYDILTGSWFGAFDI | IGKV1-17 | IGKJ4 | LQHNSYPLT |
| COV2-2210 | IGHV3-30-3 | IGHJ4 | ARDQEWFRELFLFDY | IGKV1-12 | IGKJ3 | QQANSFPPT |
| COV2-2369 | IGHV3-30 | IGHJ4 | AKDFGGDNTAMVEYFFDF | IGKV1-5 | IGKJ1 | QQYNSYSPT |
| COV2-2490 | IGHV3-7 | IGHJ4 | ARDPYDLYGDYGGTFDY | IGKV1-5 | IGKJ4 | QQYNSYSLT |
| COV2-2582 | IGHV7-4-1 | IGHJ6 | ARDQDSGYPTYYYYYMDV | IGKV2D-29 | IGKJ4 | MQSIQPPLT |
| COV2-2660 | IGHV3-13 | IGHJ6 | ARADPYQLLGQHYYYGMDV | IGKV3-20 | IGKJ5 | QQYGSSPLIT |
| DH1052 | IGHV1-69-2 | IGHJ4 | ATSSGPSRLCGGGSCYHSFDY | IGKV3-20 | IGKJ1 | QQYGSSPTWT |
| DH1053 | IGHV3-43 | IGHJ4 | AKAKDPYTEYFDY | IGKV1-5 | IGKJ4 | QQYYIYSLS |
| DH1054 | IGHV3-53 | IGHJ3 | ARGDIVGATWDPAFDI | IGLV3-21 | IGLJ2 | QVWDTSSDHSV V |
| DH1055 | IGHV2-70 | IGHJ4 | ARINAYSSSWPTFDY | IGKV3-20 | IGKJ1 | QQYGSSSWT |
| DH1056 | IGHV4-39 | IGHJ5 | ARSSSGFSYDTPLDP | IGKV1-17 | IGKJ5 | LQHNSYPIT |

**Table S2. Healthy donor BCR repertoire sequences obtained from public databases.**

| Status | Study | Donors | Clones | Reference |
|---|---|---|---|---|
| Healthy | Meng-2017 | 6 | 1493039 | (Meng et al., 2017) |
| | Setliff-2018 | 6 | 130109 | (Setliff et al., 2018) |
| | Gidoni-2019 | 97 | 1689815 | (Gidoni et al., 2019) |
| | Soto-2019 | 3 | 972608 | (Soto et al., 2019) |
| | Turner-2020 | 3 | 237411 | (Turner et al., 2020) |
| | Ghraichy-2020 | 53 | 1932769 | (Ghraichy et al., 2020) |
| | Roskin-2020 | 114 | 48313781 | (Roskin et al., 2020) |
| | Zhang-2020 | 5 | 2359 | (Zhang et al., 2020) |
| | Wen-2020 | 4 | 1968 | (Wen et al., 2020) |
| | Zhou-2021 | 6 | 627470 | (Zhou et al., 2021) |
| | Total | 297 | 55401329 | |

**Table S3. COVID-19 patient BCR repertoire sequences obtained from public databases.**

| Status | Study | Donors | Clones | Reference |
|---|---|---|---|---|
| COVID-19 | Niu-2020 | 11 | 485034 | (Niu et al., 2020) |
| | Nielsen-2020 | 5 | 155297 | (Nielsen et al., 2020) |
| | Galson-2020 | 19 | 745666 | (Galson et al., 2020) |
| | Montague-2020 | 19 | 72148 | (Montague et al., 2021) |
| | Schultheiß-2020 | 37 | 264906 | (Schultheiss et al., 2020) |
| | Cervantes-2020 | 10 | 121728 | (Kuri-Cervantes et al., 2020) |
| | Wen-2020 | 10 | 6207 | (Wen et al., 2020) |
| | Zhang-2020 | 13 | 8229 | (Zhang et al., 2020) |
| | Bernardes-2020 | 13 | 145323 | (Bernardes et al., 2020) |
| | Woodruff-2020 | 2 | 5476 | (Woodroof et al., 2020) |
| | Kim-2021 | 17 | 4984993 | (Kim et al., 2021) |
| | Mor-2021 | 14 | 70122 | (Mor et al., 2021) |
| | Zhou-2021 | 30 | 1295077 | (Zhou et al., 2021) |
| | Goel-2021 | 5 | 86853 | (Goel et al., 2021) |
| | Sokal-2021 | 8 | 43594 | (Sokal et al., 2021) |
| | Total | 213 | 8490653 | |

**Table S4. BNT162b2 vaccinated donor BCR repertoire sequences obtained from public databases.**

| Status | Study | Donors | Clones | Reference |
|---|---|---|---|---|
| BNT162b2 | Goel-2021-2 | 4 | 185543 | (Goel, Painter, et al., 2021) |
| vaccinated | Schmitz-2021 | 22 | 195252 | (Schmitz et al., 2021) |
| | Turner-2021 | 3 | 10406 | (Turner et al., 2021) |
| | Total | 29 | 391201 | |
